# A naturally occurring variant of *MBD4* causes maternal germline hypermutation in primates

**DOI:** 10.1101/2023.03.27.534460

**Authors:** Alexandra M. Stendahl, Rashesh Sanghvi, Samuel Peterson, Karina Ray, Ana C. Lima, Raheleh Rahbari, Donald F. Conrad

**Author notes:** These authors contributed equally (Alexandra M. Stendahl, Rashesh Sanghvi, Samuel Peterson). These authors jointly supervised the project (Raheleh Rahbari, Donald F. Conrad).

## Abstract

As part of an ongoing genome sequencing project at the Oregon National Primate Research Center, we identified a rhesus macaque with a rare homozygous frameshift mutation in the gene Methyl-CpG binding domain 4 (*MBD4*). MBD4 is responsible for the repair of C>T deamination mutations at CpG locations and has been linked to somatic hypermutation and cancer predisposition in humans. We show here that MBD4-associated hypermutation also affects the germline: the 6 offspring of the *MBD4*-null dam have a 4-6 fold increase in *de novo* mutation burden. This excess burden was predominantly C>T mutations at CpG locations consistent with *MBD4* loss-of-function in the dam. There was also a significant excess of C>T at CpA sites, indicating an important, underappreciated role for MBD4 to repair deamination in CpA contexts. The *MBD4*-null dam developed sustained eosinophilia later in life, but we saw no other signs of neoplastic processes associated with *MBD4* loss-of-function in humans, nor any obvious disease in the hypermutated offspring. This work provides what is likely the first evidence for a genetic factor causing hypermutation in the maternal germline of a mammal, and adds to the very small list of naturally occurring variants known to modulate germline mutation rates in mammals.

## Main Text

The DNA glycosylase *MBD4* (Methyl-CpG binding domain 4, MIM: 603574) repairs C>T mutations through the base excision repair pathway by removing the thymine in a G-T mismatch.^1^ *MBD4* is specifically active at repairing the spontaneous deamination of 5-methyl-cytosine to thymine, one of the most common somatic mutations in the genome.^2^ MBD4 is critical for genome stability and preventing mutations, and loss of function of *MBD4* has recently been associated with higher risk of MBD4-associated neoplasia syndrome (MIM: 619975), a multi-organ tumor predisposition syndrome ^3,4^. It is further associated with several specific cancers including adenomatous colorectal polyposis, acute myeloid leukemia, and uveal melanoma ^3–5^. Additionally, many types of cancerous tumors are often found to contain mutations in the *MBD4* gene, especially gastric, endometrial, and pancreatic carcinomas ^6^. Congruent with the function of *MBD4*, C>T mutations are found in high incidence in gastro-intetinal tumors of *MBD4*^-/-^ mice, with most occurring at CpG locations ^6^.

As part of an ongoing genetic study at Oregon National Primate Research Center (ONPRC), we identified a rhesus macaque (ID: 26537) with a germline homozygous TC>C deletion in the *MBD4* gene at 2:147,059,371 (MMul 10 genome assembly). Sequencing of the parents of 26537 confirmed that the *MBD4* genotype is an inherited homozygous deletion (**Figure 1)**. In macaques, the *MBD4* protein is 572 amino acids long and consists of a methyl binding domain at the N terminus and a glycosylase domain at the C terminus, which is involved in the DNA repair. The mutation in 26537 results in a frameshift mutation (Gly329fs) leading to a isoleucine to serine substitution at position 330 (Ile330Ser) and an early truncation at the following amino acid (Ile331AUU). Monkey 26537 is the only known homozygote at ONPRC or in mGAP, the database of macaque genomes (currently 2,425 monkeys, ^7^). The allele frequency in mGAP is 0.0072, with 30 heterozygotes in the population at ONPRC.

**Figure 1.**
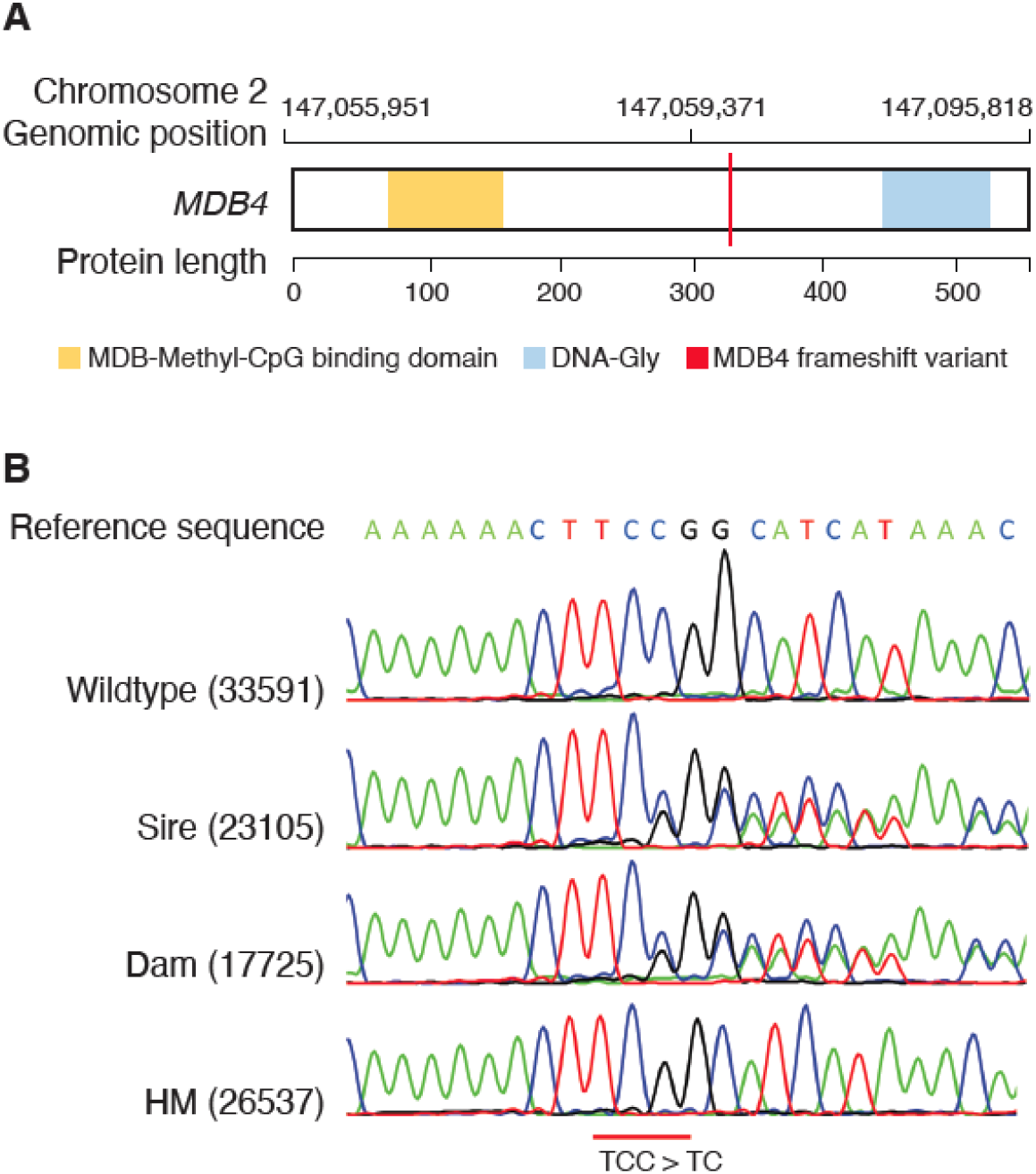
Identification of a biallelic frameshift mutation in MBD4. (A) Location of the mutation with respect to genome and protein coordinates. (B) Sanger validation of the frameshift indicates germline inheritance from both parents of 26537. HM= hypermutator. Sire and Dam are parents of 26537.

26537 was an Indian-origin rhesus macaque born at ONPRC in 2007. She gave birth to six offspring between the ages of 4 and 10, and was euthanized at age 14. Through this larger genetic study, we sequenced 26537’s six offspring, and the sires of her offspring to obtain complete trios (**Figure 2, Table S1**). We also sequenced a large pedigree of macaques from a prolific male breeder (ID: 18607), including his 111 offspring and the 61 dams (**Table S1**). Three of 26537’s offspring were sired by this prolific male breeder.

**Figure 2.**
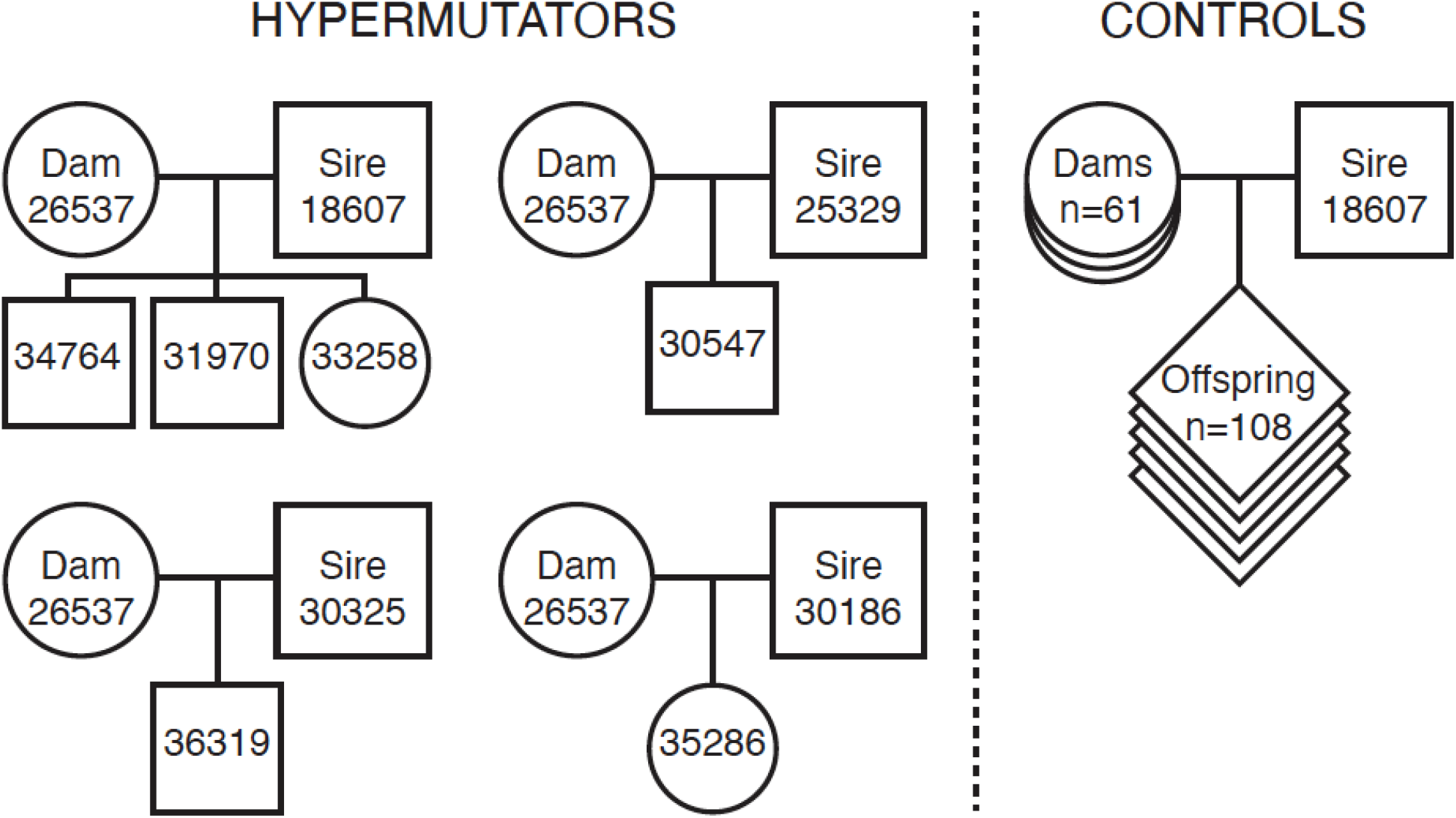
Overview of families sequenced for the study.

In our analysis of the prolific male’s pedigree, all of 26537’s offspring were flagged due to the high number of *de novo* mutations. Germline *de novo* mutations are mutations which appear in an offspring without detection in the somatic cells of either parent. Humans have a germline mutation rate of 1.2 to 1.3 × 10^-8^ per position per generation (~60-70 *de novo* mutations), while macaques appear to have a mutation rate of 0.58 to 0.77 × 10^-8^ per site per generation ^8–12^. Germline *de novo* mutations can originate as errors in DNA replication during gametogenesis or other processes. Males tend to pass on more *de novo* mutations to their offspring, which may be a result of the continuous germ cell division in spermatogenesis as opposed to oogenesis in females, which is finished before birth ^13^. However, as Abascal et al. found that somatic mutation rate is more related to tissue type than number of divisions, there may be additional mutational processes involved in *de novo* mutations in sperm^14^. Older males are especially at risk of passing high numbers of mutations to their offspring, contributing approximately 1.5-2 mutations per year of age ^9,10,14,15^. While some studies do show a maternal age effect, it is considerably smaller, approximately 0.3-0.5 mutations per year ^9,16,17^.

While the average rhesus macaque in this larger study carried 10.5 *de novo* mutations (stdev=4), offspring of 26537 carried 46-64 (**Table S1**, **Figure 3A–3B**). Despite being sired by 4 different males, all six offspring of 26537 had extremely high numbers of C>T mutations at CpG locations (between 25-39) compared to the average of 1 mutation per monkey not birthed by 26537. Because *MBD4* repairs C>T mutations at CpG locations, an enrichment in C>T mutations at CpG locations originating from the maternal haplotype is consistent with the *MBD4* mutation. Moreover, read-backed phasing revealed a 16-23 fold increase in maternally-derived mutations in offspring of 26537 compared to control trios (**Figure 3C**, **Table S2**, average of 21.33 maternal mutations vs 1.43, p=1 x 10^-95^ Poisson regression), while there was no difference in paternally-derived mutations (5.0 vs 4.1, p=0.176).

**Figure 3.**
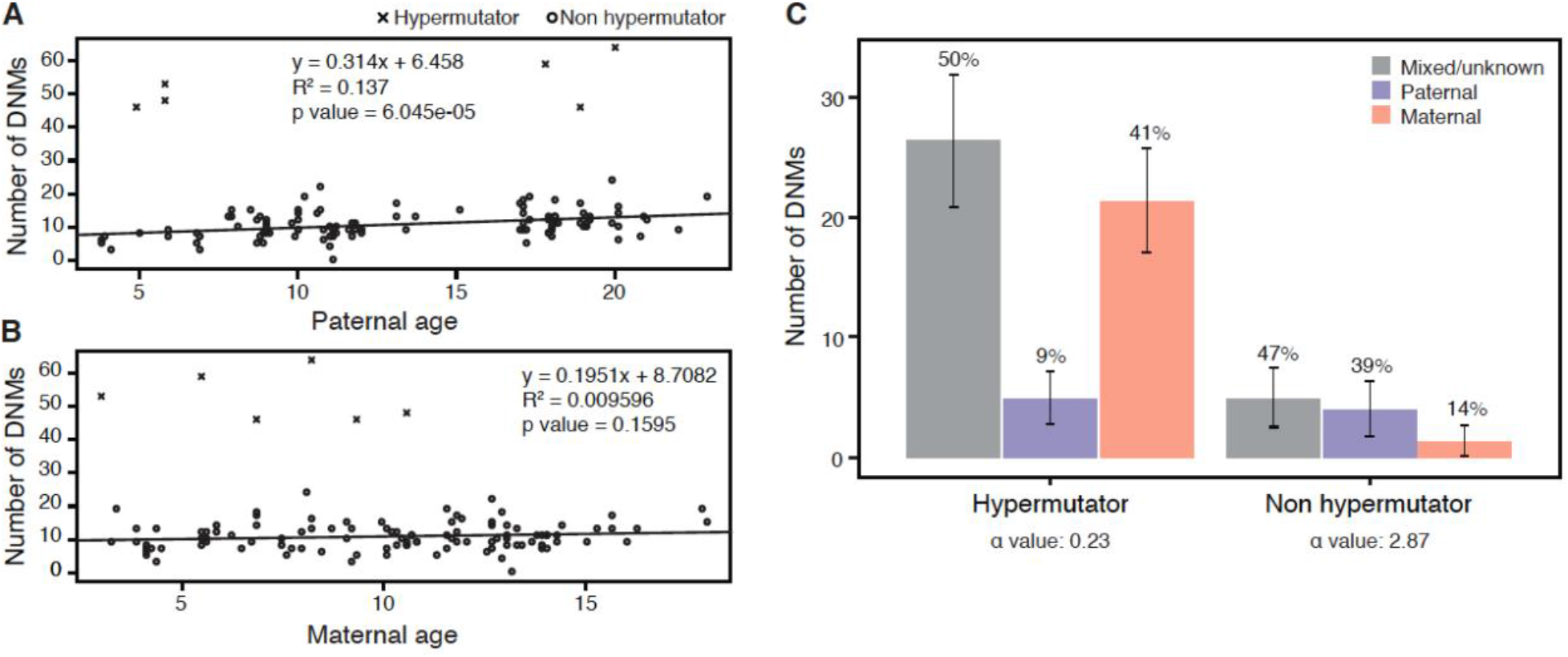
Children of 26537 show rates of germline *de novo* mutation 4-5 higher than children from control parents. (A) relationship between paternal age and number of DNMs observed in offspring, (B) relationship between maternal age and number of DNMs observed in offspring, (C) Percentage of de novo mutations observed on paternally and maternally inherited chromosomes. Read-based phasing of DNMs shows an excess of maternally-derived DNMs in offspring of 26537 not observed in control offspring. The alpha estimate for each set of phased mutations is shown. Alpha=ratio of paternal:maternal mutations.

To build further evidence implicating *MBD4* loss-of-function as the cause of the observed hypermutation, we screened 26537 for other potentially causal mutations in genes potentially related to genome instability genes (**Tables S3-S6**), assessing all other parents included in our study to provide context. These results confirmed that *MBD4* is the most likely cause of hypermutation (**Figure 4**).

**Figure 4.**
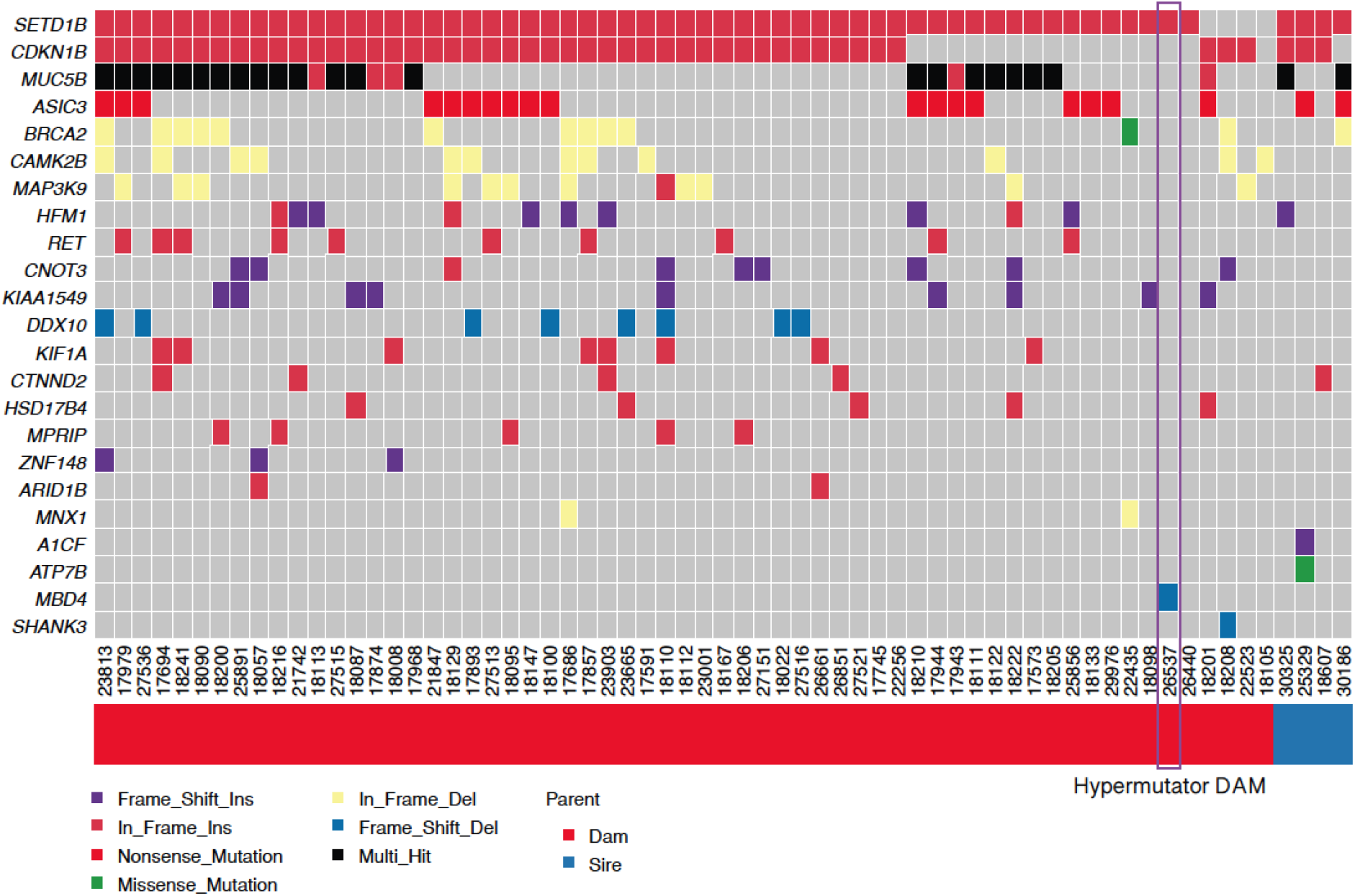
Assessment of damaging variation in candidate hypermutator genes across all parents in the study. We searched for biallelic damaging mutations across 1,381 candidate genes for germline hypermutation derived from the literature (**Table S3**). We identified candidate mutations in 23 genes, shown here. *MBD4* was the only gene specifically mutated in 26537 and not other parents in the study.

Mutational signature analysis ^18^ of all 26537 offsprings showed almost all the mutations were attributed to signature 1, while in the control trios only 20% of mutations were attributed to signature 1 and the rest (80%) were signature 5 (**Figure 5A**). This composition of mutation signatures is consistent with the mutation data in humans: somatic mutations from *MBD4-/-* individuals have an excess of signature 1 ^3^, while germline mutations in typical humans are a mixture of signatures 1 and 5 ^15^. Mutation signature 1 has been attributed to cytosine deamination, and is primarily composed of C>T mutations. Indeed, the vast majority of mutational excess observed in the offspring of 26537 were C>T mutations (**Figure 5B**). While most cytosine deamination is thought to occur at CpGs, approximately 40% of the C>T mutations in hypermutator offspring were not in CpG. We used poisson regression to test for enrichment of C>T mutations across trinucleotide contexts, and observed a clear enrichment in three trinucleotides that did not contain CpG: ACA, CCA, GCA (**Figure 5C**). This indicates a clear pattern of C>T mutations in locations where a cytosine is followed by an adenine. While CpG methylation is maintained in all tissues over the lifespan of an animal, CpA methylation is observed primarily in fetal development and in adult neurons ^19^.

**Figure 5.**
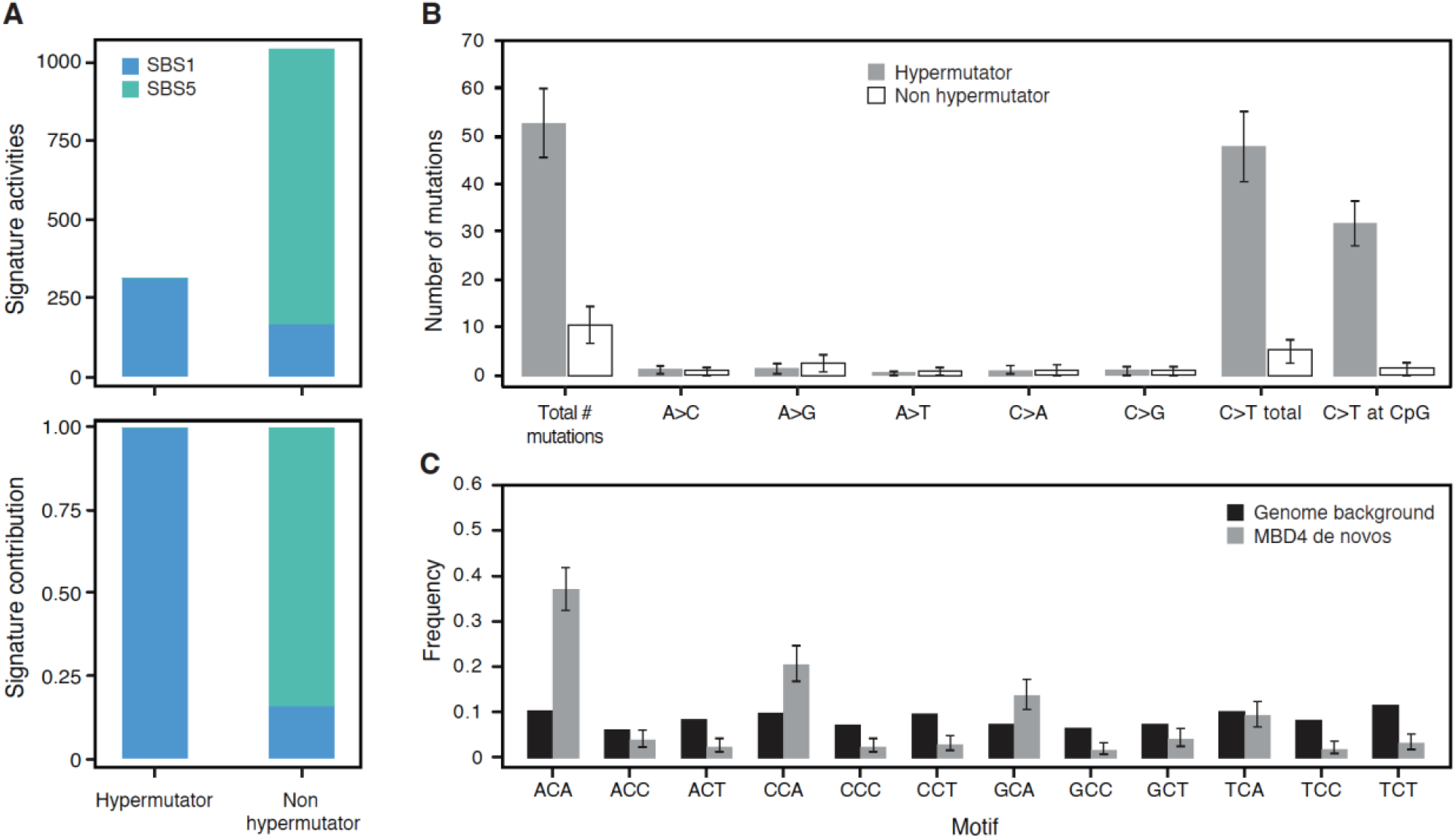
Mutation signature analysis of MBD4 germline mutations. (A) Unlike germline mutational signatures in control individuals, which are mostly signature 5 with smaller proportion of signature 1, the majority of mutations in the hypermutated offspring were attributed to signature 1 with very small contribution from signature 5. (B) Of the 6 possible (folded) substitution types, C>T was the only based substitution with a significant excess in offspring of 26537 (Hypermutator). Most of these C>T changes were in a CpG context, consistent with the known function of MBD4 in repairing spontaneous methyl-cytosine deamination. (C) The remaining excess of C>T changes were observed at CpA positions.

In order to understand whether the hypermutation phenotype or *MBD4* frameshift mutation had an effect on offspring or dam health, we reviewed electronic health records and necropsy information if present for these individuals. None of the hypermutated offspring appear to have any major health concerns. While alive, 26537 did not appear to exhibit any major health issues, aside from a single stillbirth, however tissue from the stillborn was not available for analysis. 26537 was necropsied at 14 years of age. There were no gross indications of neoplasia at necropsy, nor during histological analysis of brain, gastric tissues or lymph nodes. However, four blood chemistry tests (complete blood counts; CBCs) run on 26537 during the last year of her life revealed sustained eosinophilia, with an average measurement of 19.8% eosinophils. For context, we compared these measures to 24,638 CBCs run on 4,699 unique animals at ONPRC since 2016. The average % EOS for animals tested over this time was 2.2%; only 71 tests (0.3%) reported higher EOS counts (**Figure 6**). Fecal parasitology excluded parasitemia as a possible cause for eosinophilia in 26537. Eosinophilia is sometimes observed in several cancers and cancer-related syndromes, especially eosinophilic leukemia and myelodysplastic syndrome (MDS). Both MDS and acute myeloid leukemia have been observed in humans with biallelic mutation of *MBD4* ^3^. These findings provide evidence that the MBD4 mutations in 26537 potentially had functional consequences at a cellular level, beyond deficiencies in DNA repair.

**Figure 6.**
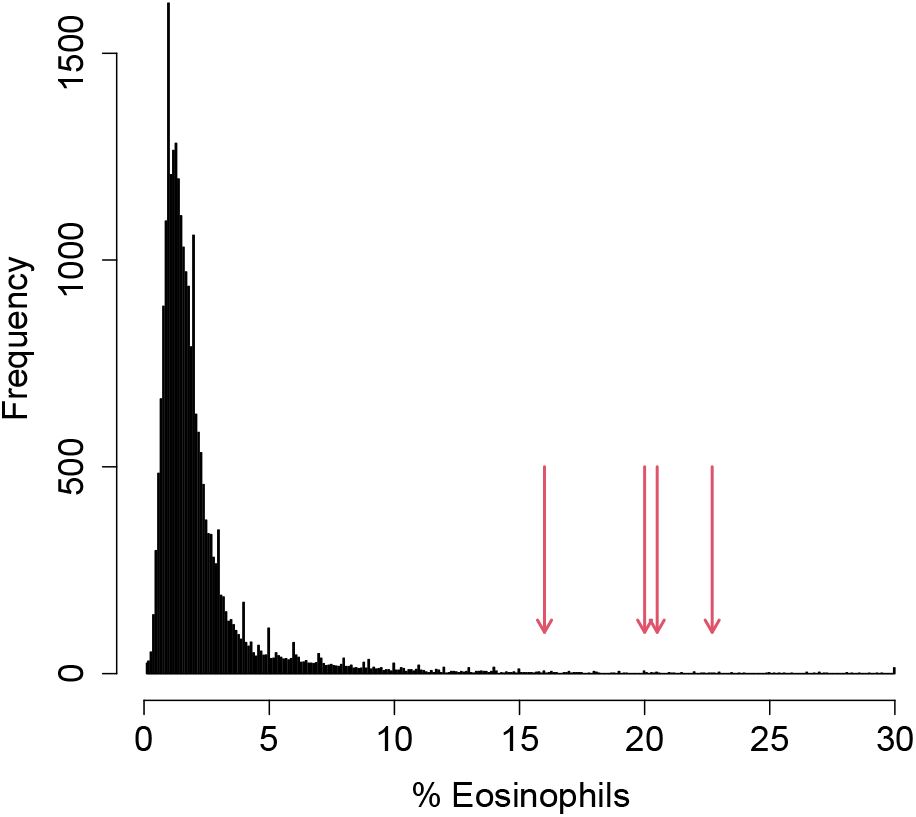
26537 showed sustained eosinophilia at age 14. Four independent CBCs run over several weeks showed 16%-22.7% eosinophils (red arrows). These numbers were in the top 0.5% of over 25,000 CBCs run at ONPRC since 2016 (histogram).

In conclusion, through a large genetic sequencing project of macaques at ONPRC, six hypermutated monkeys were identified, all originating from a dam with a homozygous mutation in the *MBD4* gene causing a premature stop codon. This discovery adds to the short list of naturally occurring genetic variants that have been found to modulate germline mutation rates, in humans, in mice, and now in rhesus macaques.

Hypermutation can be the result of both environmental factors, such as parental exposure to chemotherapy, as well as genetic factors, such as a knockout of a DNA repair gene. Germline hypermutation (a large excess of *de novo* mutations) is quite rare in humans. Kaplanis et al. found twelve individuals out of 21,879 human families with a hypermutated phenotype; only 2 of these could be attributed to genetic causes ^20^. Both cases were born to fathers with a germline hypermutation phenotype, attributed to rare biallelic mutations in the DNA repair genes *XPC* (MIM: 613208) and *MPG* (MIM: 156565). *MPG* encodes N-methylpurine DNA glycosylase, which, similar to *MBD4*, is involved in correction of deaminated purines by the base-excision repair pathway. Inherited variation in another DNA glycosylase, *MUTYH*, can increase somatic mutation rates in humans ^21^, and in the germline in mice ^22^. Notably, our finding of a maternal germline hypermutator phenotype appears to be the first clear evidence of genetic variants affecting germline mutation in a female mammal. Because the baseline mutation rate in male primates (including humans) is much higher than in females, we expect that statistical power will continue to favor ascertainment of male germline hypermutators when studying population samples, even for genetic variants that affect male and female germlines equally.

Many questions remain about how, and when during the lifecycle, the germline is vulnerable to DNA damage. The data presented here may hold some important clues regarding the timing of cytosine deamination and its repair in the maternal germline. The *de novo* mutations transmitted from 26537 to her offspring were present in the oocytes that generated each offspring. All offspring from 26537 were heterozygous for the MBD4 mutation, meaning that they were formed by fertilization of an *MBD4^-/-^* oocyte by a MBD4+ spermatozoa. In principle, the mutations in the *MBD4^-/-^* oocytes could have originated at any timepoint in the lifecycle of 26537, from the zygote to the fertilization of each *MBD4^-/-^* oocyte. Multiple observations in this study would seem to restrict the plausible developmental window during which the hypermutator phenotype was active. First, there is no detectable effect of maternal age on the number of mutations in the offspring of 26537 (**Figure 3B**). This implies that the excess germline mutation burden in each oocyte from 26537 is not an accumulation of spontaneous deamination over the lifetime of the animal. Second, there is minimal sharing of mutations among the offspring of 26537 - 4 of 316 (1.2%) of filtered mutations in the offspring were observed more than once (**Table S7**). This suggests that none of the mutations were present in the embryonic progenitor of all germ cells, and very few of the 316 mutations were present in the embryo at the time of the PGC specification. Finally, in addition to the expected excess of CpG mutations in the offspring of 26537, we observed a secondary enrichment in CpA contexts accounting for 28% of all calls in these children (**Figure 5C**). non-CpG methylation, especially involving CpA, occurs genomewide in oocytes of mice ^23^ and human ^24^. Human oocytes show dramatic (>100%) increases in CpA methylation during oocyte maturation, while CpG methylation levels stay stable ^24^. Furthermore, *MBD4* is expressed in developing oocytes, and its expression is highly correlated to the oocyte expression levels of *DNMT3A* (MIM:602769), a methyltransferase responsible for non-CpG methylation. Both *MBD4* and *DNMT3A* show a remarkable increase in transcription in human fetal ovary during 16 weeks gestation, when oocytes are first reaching the germinal vesicle stage ^25^. Taken together, we propose that the source of the germline hypermutation phenotype from *MBD4-/-* animals is not due to a loss of “maintenance” deamination repair that accumulates over the lifetime of the animal, but, instead due to a time-restricted window of damage, perhaps induced during *de novo* CpA methylation.

## Data and code availability

The raw whole-genome sequencing data for this study has been uploaded to SRA under bioproject number PRJNA382404. Mapping of sample-level IDs to SRA accessions and mGAP IDs are provided in **Table S8**.

De Novo Gear: https://github.com/ultimatesource/denovogear#dng-dnm-finding-denovo-mutations-in-trios-and-pairs

PhaseMyDeNovo: https://github.com/queenjobo/PhaseMyDeNovo

SigProfileExtractor: https://github.com/AlexandrovLab/SigProfilerExtractor

## Supplemental Information

Supplemental Information can be found online at XXXX

## Acknowledgements

We thank Rhonda MacAllister and Carolyn Labriola for help with pathology and medical records, and Katinka Vigh-Conrad for assistance with figures. D.F.C. is supported by National Institute of Health Office of Directors (NIH/OD) Grant P51 OD011092 (to the Oregon National Primate Research Center).

## Web Resources

MGAP database, http://mgap.ohsu.edu

## Supplementary Table Captions

Table S1: Sample Information

Table S2: Phased Variants by PhaseMyDeNovo

Table S3: Candidate genes for hypermutator phenotype

Table S4. High and Moderate impact variants within the genes of interest (GOI) genes. Impact is determined based on VEP annotation.

Table S5: High and Moderate impact homozygous variants within the genes of interest (GOI) genes. Impact is determined based on VEP annotation.

Table S6: High and Moderate impact homozygous variants within the top 100 mutated genes of interest (GOI) genes. Impact is determined based on VEP annotation.

Table S7: Mutations observed in multiple offspring of 26537.

Table S8: SRA and MGaP ID mapping for samples in the study.

## References

1. Pidugu, L.S., Bright, H., Lin, W.-J., Majumdar, C., Van Ostrand, R.P., David, S.S., Pozharski, E., and Drohat, A.C. (2021). Structural Insights into the Mechanism of Base Excision by MBD4. J. Mol. Biol. 433, 167097.

2. Sanders, M.A., Chew, E., Flensburg, C., Zeilemaker, A., Miller, S.E., Al Hinai, A.S., Bajel, A., Luiken, B., Rijken, M., Mclennan, T., et al. (2018). MBD4 guards against methylation damage and germ line deficiency predisposes to clonal hematopoiesis and early-onset AML. Blood 132, 1526–1534.

3. Palles, C., West, H.D., Chew, E., Galavotti, S., Flensburg, C., Grolleman, J.E., Jansen, E.A.M., Curley, H., Chegwidden, L., Arbe-Barnes, E.H., et al. (2022). Germline MBD4 deficiency causes a multi-tumor predisposition syndrome. Am. J. Hum. Genet. 109, 953–960.

4. Derrien, A.-C., Rodrigues, M., Eeckhoutte, A., Dayot, S., Houy, A., Mobuchon, L., Gardrat, S., Lequin, D., Ballet, S., Pierron, G., et al. (2021). Germline MBD4 Mutations and Predisposition to Uveal Melanoma. J. Natl. Cancer Inst. 113, 80–87.

5. Millar, C.B., Guy, J., Sansom, O.J., Selfridge, J., MacDougall, E., Hendrich, B., Keightley, P.D., Bishop, S.M., Clarke, A.R., and Bird, A. (2002). Enhanced CpG mutability and tumorigenesis in MBD4-deficient mice. Science 297, 403–405.

6. Wong, E., Yang, K., Kuraguchi, M., Werling, U., Avdievich, E., Fan, K., Fazzari, M., Jin, B., Brown, A.M.C., Lipkin, M., et al. (2002). Mbd4 inactivation increases Cright-arrowT transition mutations and promotes gastrointestinal tumor formation. Proc. Natl. Acad. Sci. U. S. A. 99, 14937–14942.

7. Bimber, B.N., Yan, M.Y., Peterson, S.M., and Ferguson, B. (2019). mGAP: the macaque genotype and phenotype resource, a framework for accessing and interpreting macaque variant data, and identifying new models of human disease. BMC Genomics 20, 176.

8. Conrad, D.F., Keebler, J.E.M., DePristo, M.A., Lindsay, S.J., Zhang, Y., Casals, F., Idaghdour, Y., Hartl, C.L., Torroja, C., Garimella, K.V., et al. (2011). Variation in genome-wide mutation rates within and between human families. Nat. Genet. 43, 712–714.

9. Kong, A., Frigge, M.L., Masson, G., Besenbacher, S., Sulem, P., Magnusson, G., Gudjonsson, S.A., Sigurdsson, A., Jonasdottir, A., Jonasdottir, A., et al. (2012). Rate of de novo mutations and the importance of father’s age to disease risk. Nature 488, 471–475.

10. Rahbari, R., UK10K Consortium, Wuster, A., Lindsay, S.J., Hardwick, R.J., Alexandrov, L.B., Al Turki, S., Dominiczak, A., Morris, A., Porteous, D., et al. (2016). Timing, rates and spectra of human germline mutation. Nat. Genet. 48, 126–133.

11. Wang, R.J., Thomas, G.W.C., Raveendran, M., Harris, R.A., Doddapaneni, H., Muzny, D.M., Capitanio, J.P., Radivojac, P., Rogers, J., and Hahn, M.W. (2020). Paternal age in rhesus macaques is positively associated with germline mutation accumulation but not with measures of offspring sociability. Genome Res. 30, 826–834.

12. Bergeron, L.A., Besenbacher, S., Bakker, J., Zheng, J., Li, P., Pacheco, G., Sinding, M.-H.S., Kamilari, M., Gilbert, M.T.P., Schierup, M.H., et al. (2021). The germline mutational process in rhesus macaque and its implications for phylogenetic dating. Gigascience 10,.

13. Crow, J.F. (2000). The origins, patterns and implications of human spontaneous mutation. Nat. Rev. Genet. 1, 40–47.

14. Abascal, F., Harvey, L.M.R., Mitchell, E., Lawson, A.R.J., Lensing, S.V., Ellis, P., Russell, A.J.C., Alcantara, R.E., Baez-Ortega, A., Wang, Y., et al. (2021). Somatic mutation landscapes at single-molecule resolution. Nature 593, 405–410.

15. Moore, L., Cagan, A., Coorens, T.H.H., Neville, M.D.C., Sanghvi, R., Sanders, M.A., Oliver, T.R.W., Leongamornlert, D., Ellis, P., Noorani, A., et al. (2021). The mutational landscape of human somatic and germline cells. Nature 597, 381–386.

16. Jónsson, H., Sulem, P., Kehr, B., Kristmundsdottir, S., Zink, F., Hjartarson, E., Hardarson, M.T., Hjorleifsson, K.E., Eggertsson, H.P., Gudjonsson, S.A., et al. (2017). Parental influence on human germline de novo mutations in 1,548 trios from Iceland. Nature 549, 519–522.

17. Deciphering Developmental Disorders Study (2017). Prevalence and architecture of de novo mutations in developmental disorders. Nature 542, 433–438.

18. Tate, J.G., Bamford, S., Jubb, H.C., Sondka, Z., Beare, D.M., Bindal, N., Boutselakis, H., Cole, C.G., Creatore, C., Dawson, E., et al. (2019). COSMIC: The Catalogue Of Somatic Mutations In Cancer. Nucleic Acids Res. 47, D941–D947.

19. Greenberg, M.V.C., and Bourc’his, D. (2019). The diverse roles of DNA methylation in mammalian development and disease. Nat. Rev. Mol. Cell Biol. 20, 590–607.

20. Kaplanis, J., Ide, B., Sanghvi, R., Neville, M., Danecek, P., Coorens, T., Prigmore, E., Short, P., Gallone, G., McRae, J., et al. (2022). Genetic and chemotherapeutic influences on germline hypermutation. Nature 605, 503–508.

21. Robinson, P.S., Thomas, L.E., Abascal, F., Jung, H., Harvey, L.M.R., West, H.D., Olafsson, S., Lee, B.C.H., Coorens, T.H.H., Lee-Six, H., et al. (2022). Inherited MUTYH mutations cause elevated somatic mutation rates and distinctive mutational signatures in normal human cells. Nat. Commun. 13, 3949.

22. Sasani, T.A., Ashbrook, D.G., Beichman, A.C., Lu, L., Palmer, A.A., Williams, R.W., Pritchard, J.K., and Harris, K. A natural mutator allele shapes mutation spectrum variation in mice.

23. Shirane, K., Toh, H., Kobayashi, H., Miura, F., Chiba, H., Ito, T., Kono, T., and Sasaki, H. (2013). Mouse oocyte methylomes at base resolution reveal genome-wide accumulation of non-CpG methylation and role of DNA methyltransferases. PLoS Genet. 9, e1003439.

24. Yu, B., Dong, X., Gravina, S., Kartal, Ö., Schimmel, T., Cohen, J., Tortoriello, D., Zody, R., Hawkins, R.D., and Vijg, J. (2017). Genome-wide, Single-Cell DNA Methylomics Reveals Increased Non-CpG Methylation during Human Oocyte Maturation. Stem Cell Reports 9, 397–407.

25. Galetzka, D., Weis, E., Tralau, T., Seidmann, L., and Haaf, T. (2007). Sex-specific windows for high mRNA expression of DNA methyltransferases 1 and 3A and methyl-CpG-binding domain proteins 2 and 4 in human fetal gonads. Mol. Reprod. Dev. 74, 233–241.

