## Supplementary Information for "A naturally occurring variant of *MBD4* causes maternal germline hypermutation in primates"

### Supplementary Methods Stendahl, Sanghvi, Peterson et al.

Initial candidate *de novo* mutations were identified using DeNovoGear^1^. We selected sites with a probability of *de novo* single base pair mutation greater than 0.5. These sites were further filtered to include sites where offspring and parent read depth was between 4 and 80 reads. Offspring must have a variant allele frequency (VAF) between 35% and 70% for mutations on autosomes. Mutations on the X chromosome should have a VAF between 35% and 70% for female offspring and greater than 70% for male offspring. Mutations on the Y chromosome were excluded. Alternate allele reads must appear on both forward and reverse strands, with at least 2 reads supporting the alternate allele. We removed sites which fell in highly repetitive regions (UCSC simple repeats track) and in known segmental duplications (UCSC Segmental Duplication track)^2^. We removed variants where either parent had more than one alternate allele read or greater than 10% variant allele frequency. We removed mutations which were relatively common in the macaque population, as defined by having an allele frequency in mGAP greater than 0.01. Finally, we used a binomial test with FDR correction (adjusted p-value < 0.05) to settle on our final set of *de novo* mutations.

Because paternal age is a major factor in *de novo* mutation counts, we analyzed the contribution of both paternal and maternal age to mutation counts. We used poisson regression to model the effects of parental age, hypermutator status, and sequencing coverage on observed mutation counts. For the non-hypermutated group, the number of *de novo* mutations was correlated to paternal age (p = 6.04 x 10-5) and increased by approximately 0.31 mutations per year as the sire aged (y=0.31x + 6.5). Maternal age showed no increase with maternal age (Y= 0.20x - 8.7, p = 0.15). Hypermutated individuals were sired by males ranging in age from 4.9 - 20.

To determine the mutational processes resulting in the hypermutation in the six offspring of 26537, we compared the *de novo* mutation profiles by extracting mutational signatures using SigProfilerExtractor (v 1.1.7) and estimating their contribution to the mutation burden.

To confirm the hypermutation phenotype was a result of mutations coming from 26537, we phased mutations to either the maternal or paternal allele. Germline mutations were jointly called for each trio using HaplotypeCaller ^3^ and the *de novo* mutations were phased using PhaseMyDeNovo (<https://github.com/queenjobo/PhaseMyDeNovo>). We were able to phase 50% of mutations to a single parent in hypermutators and 53% of mutations in non-hypermutators. Non-hypermutated animals had 74% of phaseable mutations phased to the paternal side, whereas the hypermutated offspring had 81% of phaseable mutations phased to the maternal side. The paternal-to-maternal ratio (𝛼 value = 2.87) of the non-hypermutators was similar to that reported in literature (~3) ^4^, compared to 0.23 for hypermutators.

To determine if this loss of function in the MBD4 gene was responsible for the hypermutated phenotype, we compared the mutated genes from the six offspring of 26537 to the rest of the pedigree. We annotated the *de novo* mutations using VEP and investigated homozygous mutations in genes from DNA damage repair pathways (n=276), developmental disorder genes (n=395), cancer gene census and cancer predisposition (n=565). (**Figure 4; Supplementary Table 3**)

### Supplementary References

1. Ramu, A., Noordam, M.J., Schwartz, R.S., Wuster, A., Hurles, M.E., Cartwright, R.A., and Conrad, D.F. (2013). DeNovoGear: de novo indel and point mutation discovery and phasing. Nat. Methods *10*, 985–987.

2. Haeussler, M., Zweig, A.S., Tyner, C., Speir, M.L., Rosenbloom, K.R., Raney, B.J., Lee, C.M., Lee, B.T., Hinrichs, A.S., Gonzalez, J.N., et al. (2019). The UCSC Genome Browser database: 2019 update. Nucleic Acids Res. *47*, D853–D858.

3. Poplin, R., Ruano-Rubio, V., DePristo, M.A., Fennell, T.J., Carneiro, M.O., Van der Auwera, G.A., Kling, D.E., Gauthier, L.D., Levy-Moonshine, A., Roazen, D., et al. Scaling accurate genetic variant discovery to tens of thousands of samples.

4. Wang, R.J., Thomas, G.W.C., Raveendran, M., Harris, R.A., Doddapaneni, H., Muzny, D.M., Capitanio, J.P., Radivojac, P., Rogers, J., and Hahn, M.W. (2020). Paternal age in rhesus macaques is positively associated with germline mutation accumulation but not with measures of offspring sociability. Genome Res. *30*, 826–834.
